# Refining conformational ensembles of flexible proteins against small-angle X-ray scattering data

**DOI:** 10.1101/2021.05.29.446281

**Authors:** Francesco Pesce, Kresten Lindorff-Larsen

## Abstract

Intrinsically disordered proteins and flexible regions in multi-domain proteins display substantial conformational heterogeneity. Characterizing the conformational ensembles of these proteins in solution typically requires combining one or more biophysical techniques with computational modelling or simulations. Experimental data can either be used to assess the accuracy of a computational model or to refine the computational model to get a better agreement with the experimental data. In both cases, one generally needs a so-called forward model, i.e. an algorithm to calculate experimental observables from individual conformations or ensembles. In many cases, this involve one or more parameters that need to be set, and it is not always trivial to determine the optimal values or to understand the impact on the choice of parameters. For example, in the case of small-angle X-ray scattering (SAXS) experiments, many forward models include parameters that describe the contribution of the hydration layer and displaced solvent to the background-subtracted experimental data. Often, one also needs to fit a scale factor and a constant background for the SAXS data, but across the entire ensemble. Here, we present a protocol to dissect the effect of free-parameters on the calculated SAXS intensities, and to identify a reliable set of values. We have implemented this procedure in our Bayesian/Maximum Entropy framework for ensemble refinement, and demonstrate the results on four intrinsically disordered proteins and a three-domain protein connected by flexible linkers. Our results show that the resulting ensembles can depend on the parameters used for solvent effects, and suggests that these should be chosen carefully. We also find a set of parameters that work robustly across all proteins.

**SIGNIFICANCE:** The flexibility of a protein is often key to its biological function, yet understanding and characterizing its conformational heterogeneity is difficult. We here describe a robust protocol for combining small-angle X-ray scattering experiments with computational modelling to obtain a conformational ensemble. In particular, we focus on the contribution of protein hydration to the experiments and how this is included in modelling the data. Our resulting algorithm and software should make modelling intrinsically disordered proteins and multi-domain proteins more robust, thus aiding in understanding the relationship between protein dynamics and biological function.

## INTRODUCTION

Small-angle X-ray scattering (SAXS) experiments are widely used in the field of integrative structural biology as a versatile tool to probe conformational ensembles of biomolecules in solution. When faced with highly dynamic and flexible systems, solving a crystallographic structure may either not be possible or only provide a static image that does not capture key aspects of the system. SAXS experiments instead give a low to medium resolution ensemble-averaged view of the biomolecule. While for relatively rigid macromolecules it may be possible to derive a single averaged shape directly from a SAXS experiment (1–4), this is generally not possible for very flexible molecules. The possibility to calculate SAXS profiles from atomic coordinates, however, makes it possible to average across a distribution of conformations (a ‘prior distribution’), compare the result with an experiment and to correct the prior distribution in case of poor agreement (5, 6).

SAXS measurements report on the total scattering of X-rays from all molecules in solution. Thus, the resulting scattering profile represents the entire solution (buffer and solute). For this reason, one also collects scattering data for the buffer alone, which is then subtracted from the data from the solution with the macromolecule to get the excess scattering. As the density of solvent around the protein (the hydration layer) may differ from that of the bulk solvent (7), the resulting data (colloquially termed SAXS data, though in practice it’s a difference between two SAXS measurements) represents the signal coming from the protein together with its hydration envelope and the solvent displaced by the protein.

The calculation of SAXS intensities is done using a so-called forward model, *i.e*. an algorithm to predict an experimental observable from a structural model. Several forward models for SAXS exist and a key distinction is how they treat the scattering contribution from the protein hydration layer (5). In particular, to account for the hydration layer contribution, modelling approaches for SAXS data generally fall into two distinct categories.

In explicit solvent models (8–12), displaced water molecules and perturbed properties of the hydration envelope are explicitly taken into account in the calculation of scattering intensities. In particular, the scattering from the solute (protein) and hydration layer effects is estimated by explicit subtraction of the scattering calculated of the solvated protein and the solvent alone.

In contrast, in implicit solvent models (13–16) the contribution of the hydration layer and displaced volume to the scattering is modelled typically through one or more parameters that need to be set. The effects of the hydration layer can be modelled by a hydration shell of some width, Δ, and with excess density (and thus changed scattering compared to bulk solvent), *δ_ρ_*. Typically, only the product Δ · *δ_ρ_* is important, and thus often Δ is fixed (e.g. at 3 Å) and only *δ_ρ_* needs to be set (or fit). Similarly, implicit solvent models may include a parameter for the effective atomic radius (*r*_0_), which both affects the overall displaced volume but also to some extent the contribution of the hydration layer.

While the explicit solvent strategy may provide a more realistic representation of the hydration layer and its contribution to the scattering data, it can be computationally expensive and requires a force field and water model that accurately models protein-water interactions. While it has been shown that the calculations on folded proteins are not particularly sensitive to the choice of water model (9, 17), there is more uncertainty about the best models for protein-water interactions for disordered proteins (17–20).

In many cases, one would want to use an implicit solvent strategy to calculate scattering data because of the smaller computational overhead. On the other hand, and as described above, these methods may require setting parameters that describe e.g. the protein’s solvent envelope, and it is not always clear how best to determine these. Here we note that many forward models, including implicit-solvent-based forward models for SAXS, have mostly been developed, parametrised and benchmarked using globular and folded protein structures. For folded proteins, free-parameters in forward models may be determined using SAXS data for proteins with known structures, or fitted for a given structural model. This approach, however, is difficult to apply to disordered proteins because for these there is not a well-defined reference structure from e.g. crystallography and there is uncertainty in computational methods for generating distributions of conformations (17, 20–22), both in terms of sampling efficiently all the possible conformations with the right probabilities or in terms of parametrisation. Similarly, it is not reasonable to fit these parameters independently to each structure because of the risk of substantial overfitting (9, 17), and because it is expected that the properties of the hydration layer will not depend fundamentally on the details of the structure. Finally, a key problem is that SAXS calculations are often used to construct or bias conformational ensembles, so that a procedure needs to be able to determine the free-parameters self-consistently together with the conformational ensemble.

Here, we focus our attention on these issues with implicit solvent calculations of scattering data for heterogeneous ensembles of conformations. We illustrate the effect of varying the parameters describing the hydration layer and displaced volume on ensemble refinement of intrinsically disordered proteins (IDPs) and a flexible multi-domain protein. We do so via an iterative and self-consistent strategy to select and optimize free-parameters in SAXS calculations while at the same time constructing a conformational ensemble to represent the data.

Our approach is based on a reweighting approach that is rooted in Bayesian inference (23–31) and the Maximum-entropy principle (32–37). While these methods show similarities to other approaches based for example on genetic algorithms (6, 38, 39) or Monte Carlo processes (40, 41), they differ in how they balance prior information (often encoded in a force field) with the experimental data. This balance can be particularly important for disordered proteins where the solutions are typically severely underdetermined and where large ensembles are required to provide a realistic structural description of the conformations present in solution (38). Finally, we note that in some cases it is possible to absorb some effects of protein dynamics into the forward model rather than to represent it explicitly in the form of a conformational ensemble, and such modifications exist both for various types of NMR data (42–45) and X-ray scattering or diffraction data (46–49). While this may be useful when studying small fluctuations around a well-defined ‘average’ conformation or when the dynamics is of rigid bodies in a crystal, we here examine systems where we explicitly represent a broad conformational ensemble.

## METHODS

### Generating conformational ensembles of IDPs

We generated conformational ensembles for the polypeptide backbones of four IDPs using Flexible-meccano (50). Flexible-meccano implicitly represents a potential energy function derived from the populations of backbone dihedrals in loop regions in folded protein structures. The backbone chains are built by random sampling these potentials. Other methods exist to efficiently generate conformational ensembles of IDPs, with also the possibility of taking into account transient secondary structure elements (if part of the sequence is known to assume these) (51). We chose Flexible-meccano for most of the analyses presented here as it has been shown to generate conformational ensembles of IDPs that are in good agreement with both NMR observables and SAXS data (52–55) without the need to provide any prior knowledge about the system. As the complexity of the ensembles may be influenced by the lenght of the protein, we generated larger ensembles for the longer proteins: Hst5 (24 residues, 10000 conformers), Sic1 (90 residues, 15000 conformers), *α*-Synuclein (140 residues, 20000 conformers) and Tau (441 residues, 30000 conformers). We added side chains to the backbone structures generated by Flexible-meccano using PULCHRA (56) with default settings.

### Iterative Bayesian/Maximum-Entropy reweighting scheme

In integrative structural modelling, one approach is to use reweighting to refine probability distributions in order to improve the agreement between calculated averages and experimental values (37). Here we use the Bayesian/Maximum entropy (BME) reweighting procedure (36) that, by minimal modification of the prior distribution and taking into account the uncertainty in the experimental observable (*σ_i_*), modifies the prior weights 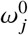 to minimise the pseudo free-energy functional (24, 26, 37):

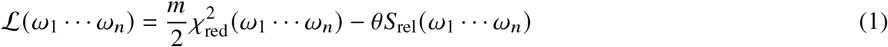

Here *m* is the number of experimental data points, (*ω*_1_… *ω_n_*) are the weights associated to each conformer of the ensemble, the reduced *χ*^2^ quantifies the agreement of the weighted average forward model predicted from each conformation *x_j_* (*F*(*x_j_*)) with the experimental data 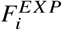:

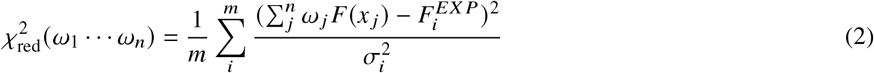

and 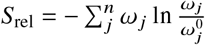 is the relative entropy that quantifies how much the reweighted distribution deviates from the prior.

Thus, when minimizing 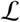 we aim to lower 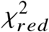 (improve agreement with experiment) while not decreasing the relative entropy term too much, in such a way to obtain the minimal modification of the prior distribution that results in a better agreement with the experimental data. The parameter *θ* is a temperature-like free-parameter that effectively sets the balance between experiments and computation, and takes into account various sources of error such as inaccuracies in the force field or the forward model (24, 37). In the limit *θ* → inf, no confidence is assigned to the experimental data and no reweighting is performed. As *θ* is decreased, more weight is put on improving agreement with experiments 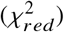, but at the cost of an increased deviation between the posterior (refined) distribution and the prior distribution (in this case generated by Flexible-meccano, or simulations with the Martini or all-atom force fields). This can also be quantified as *ϕ*_eff_ = exp(*S*_rel_), corresponding to the fraction of the original *n* frames that effectively contributes to the refined ensemble. For *θ* = 0, the *χ*^2^ is minimised without considering the prior distribution, in some cases leading to very low values of *ϕ*_eff_, and so very few conformations contribute to the final average. In methods such as BME, *θ* should be chosen in such a way to find the balance between minimising the *χ*^2^ and retaining as much information as possible from the prior, such as for example when *χ*^2^ reaches a plateau (24, 37).

We highlight that, as a consequence of the points above, the prior distribution is an important part of the procedure, as the goal of BME and related approaches is to perform the minimal modification of the prior to get a reasonable agreement with the experimental data. The definition of the prior weights is strictly dependent on the method used to sample conformations. In case of standard molecular dynamics, where the conformations are sampled from a Boltzmann distribution, or Flexible-meccano where the conformations are sampled from specific backbone dihedral angle potential wells, the weights are uniform as the probability of a certain conformation is related to its occurrence in the ensemble.

Since the goal of the BME is to decrease the *χ*^2^, it is important to ensure, when needed, that the experimental and calculated values are on the same scale. While SAXS intensities can be measured and calibrated on an absolute scale, this depends on careful calibration of the instrument and accurate measurements of the protein concentration. Thus, often calculated SAXS profiles are rescaled to match the experimental data. Moreover, experimental SAXS data may contain a small, non-zero background scattering, e.g. from imperfect background subtraction, which sometimes is dealt with by shifting calculated SAXS profiles to get a better fit.

To account for these issues, we here present the iBME (iterative Bayesian/Maximum-Entropy) approach, an iterative scheme that we have developed with the aim of coupling ensemble refinement and optimization of scale factor and constant background of the calculated SAXS profiles. The scheme is structured as follow:

1. Given a set of SAXS profiles calculated from each structure in a conformational ensemble, the corresponding ensemble-averaged SAXS profile is calculated using a set of initial (prior) weights (uniform weights in all our ensembles). We then Pesce and Lindorff-Larsen perform a weighted least squares fit between the ensemble-averaged calculated SAXS profile and the experimental SAXS profile to get slope and intercept of the resulting linear fit. Weights for the weighted least squares fit are defined as 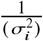.
2. The slope and intercept from (1) are used as scale factor and constant background to rescale and shift the calculated SAXS profiles.
3. BME is used for weight optimization starting from the prior weights.
4. The optimized weights from (3) are used to calculate a new ensemble average of the SAXS profiles, which in turn is used for a new weighted least square fit to the experimental profile.
5. With the new slope and intercept, the calculated SAXS dataset used in the previous BME reweighting is again rescaled and shifted.
6. Repeat 3–5 until the drop of 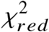 between consecutive iteration of the algorithm falls below a predefined threshold, or for a fixed number of iterations (we used 20 iterations in our analyses).

We initially tested the method using synthetic data to examine how well it can recover the scale factor and constant background (see SI text and Figs. S1–S4). We note also that iBME in part has overlap with features in BioEn (57), where only the scale factor is adjusted iteratively upon weight optimization.

iBME is implemented in an updated version of BME (https://github.com/KULL-Centre/BME). Data and scripts used for the analyses presented in this manuscript are available at https://github.com/KULL-Centre/papers/tree/main/2021/SAXS-pesce-et-al.

### Calculation of the radii of gyration

We use two different methods to estimate the average radius of gyration (*R_g_*) of a conformational ensemble, one based on the protein coordinates and another based on the SAXS data.

From a conformational ensemble, the *R_g_* for each conformer of *n* atoms can be calculated as 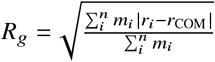, with *r_i_* being the position of the *i^th^* atom, *m_i_* its mass and 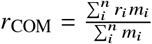 is the centre of mass. We used MDTraj (58) for these calculations, and calculate the ensemble average, 〈*R_g_*〉, as a linear or weighted average of the *R_g_* values from each conformer.

As an alternative to using the atomic masses to weigh the distances in the calculation of *R_g_*, we also use the atom contrasts, defined for each *i^th^* atom as 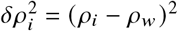, where *ρ_w_* is the density of bulk water (334 *e*/*nm*^3^) and *δ_i_* is the density of the *i^th^* atom calculated as the ratio between its number of electrons and its volume (59). We do, however, not observe substantial differences between the mass-weighted and contrast-weighted values of *R_g_* (Fig. S5).

From an experimental SAXS profile, we use the Guinier approximation to estimate the average *R_g_* in solution (60). We first transform the SAXS profile as ln *I*(*q*) vs *q*^2^, and then obtain 〈*R_g_*〉 from the slope (*a*) of a linear fit in the small-angle region using 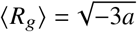. The linear fit takes into account the uncertainty on the intensities (propagated as 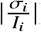) and was performed using the scikit-learn python library (61).

## RESULTS AND DISCUSSION

### Conformational ensembles and SAXS data

Our aim here is to develop a strategy to model conformational ensembles of flexible proteins with SAXS data, taking into account both uncertainty about a scale factor and constant background in the experimental SAXS data as well as effects of the hydration layer and displaced solvent. As object for our analyses we selected five proteins for which SAXS profiles had been determined experimentally and published. Also, protein flexibility may exist in multiple forms and to include different types, we first choose three IDPs of different lengths and a multi-domain protein with flexible linkers: Histatin 5 (Hst5) (SAXS data collected at 323K from (62)), Sic1 (SAXS data from (63)), full-length (ht40-)Tau (SAXS data from (64)), and the three-domain protein TIA-1 without its flexible low-complexity domain (SAXS data from (65)). Further below we also analyze an additional IDP (*α*-Synuclein, with SAXS data from (66)) to examine the robustness of the analyses done on the four proteins listed above.

We generated conformational ensembles of the four IDPs using Flexible-meccano (50). Additionally we also analyzed two previously performed molecular dynamics simulations of *α*-Synuclein (67) produced using either the Amber a99SB-*disp* or the Amber ff03ws force field. We also used a previously generated molecular dynamics simulation of TIA-1 (68). The TIA-1 simulations were performed with the Martini force field (69) after increasing the interaction strength between protein and water by 6% (68). All structures were converted to all-atom representation before calculating SAXS data.

We used the implicit solvent SAXS-calculation approach Pepsi-SAXS (14) to calculate SAXS profiles from atomic coordinates. We choose this method for its versatility and computational efficiency, but our approach will also apply to other similar methods (13, 15), and below we also discuss and show results using FoXS. When no additional information, other than atomic coordinates and an experimental SAXS profile, is provided to Pepsi-SAXS, the software may tune four parameters in order to optimize the fit between the calculated and experimental SAXS profile: (i) the intensity of the forward scattering *I*(0) (*i.e*. the scale of the profiles), (ii) a constant background *cst*, (iii) the effective atomic radius *r*_0_ and (iv) the contrast of the hydration layer *δ_ρ_*. In our calculations, we do not enable direct parameter fitting within Pepsi-SAXS and instead keep these parameters fixed to the same value for each conformer of an ensemble. As described in more detail below, we instead fit *I*(0) and *cst* as global ensemble averages, and scan *r*_0_ and *δ_ρ_* to determine self-consistent ensembles.

### Determining self-consistent ensembles and hydration layer and displaced solvent parameters

By default, Pepsi-SAXS performs a grid search for the combination of *r*_0_ and *δ_ρ_* that provides the best fit (lowest *χ*^2^) between the SAXS profile calculated from of a specific protein structure and the experimental data. While this strategy may be appropriate to calculate a SAXS profile for globular proteins with little conformational heterogeneity, it can result in overfitting if applied to each structure in a highly heterogenous conformational ensembles. Default values of *r*_0_ and *δ_ρ_* might be determined by fitting SAXS data to known crystal structures, and used without modification on other proteins. This, however, would amount to making the assumption that the hydration effects are constant and transferable from specific globular proteins to e.g. IDPs. We note here that it has been shown that the surface properties of the protein affect the hydration contribution (70, 71).

Instead, to determine the combination of parameters that best describes a conformational ensemble, to shed light on the influence of these two parameters, and to find a single set of parameters that provides a good description of the data, we here want to keep the *rationale* of a grid search, but adding an ensemble perspective. Similar to the standard grid scan, we calculate SAXS data for a range of values of *r*_0_ and *δ_ρ_*. To define the ranges for the grid, we compare the search ranges for *r*_0_ and *δ_ρ_* implemented by three of the most widely used algorithms for SAXS calculations: CRYSOL (13), FoXS (15) and Pepsi-SAXS (Table S1), and use the widest ranges allowed by the three methods. Specifically, for *r*_0_, we use 11 values in the range 1.4 Å–1.8 Å, while for *δ_ρ_* we use 30 values in the range −27.0 – 70.0 *e*/*nm*^3^. We also make the assumption that *r*_0_ and *δ_ρ_* are the same for all conformers in the ensemble. While these might in principle be conformation dependent (71, 72), we do so to decrease the risk of overfitting when varying these two parameters for each of the thousands of conformations. Also, as the goal is here to describe the conformational distribution of the protein in solution, we do not expect a substantial difference as long as there is not a strong conformational dependency on the properties of the solvation layer.

Given that the input (‘prior’) ensemble may not be fully representative of the protein in solution we do not just compare the experimental SAXS profiles with each of the average SAXS profiles calculated with Pepsi-SAXS with different values of *r*_0_ and *δ_ρ_* (37, 73). Instead, we use the BME approach to reweight the prior ensembles against the experimental data (36), using as input (one at a time) the SAXS calculations with different values of *r*_0_ and *δ_ρ_*. In the reweighting it is key that the calculated SAXS profiles match the intensity of the forward scattering *I*(0) and the constant background *cst* of the experimental signal. To accurately fit both *I*(0) and *cst* upon reweighting, we developed the iBME scheme (see Methods for detailed description and SI for validation). This new method, uses iterations of rescaling and shifting the calculated SAXS profiles and reweighting the conformational ensemble to fit a global value of *I*(0) and *cst*. The only requirement is that the same values for both *I*(0) and *cst* are used to calculate the SAXS data for all conformers (*I*(0) and *cst* are ensemble properties related to the experimental SAXS profile and independent on the single conformation). We set *I*(0) = 1 and *cst* = 0 for all conformers in the Pepsi-SAXS calculations, but as these parameters are adjusted by iBME, the choice of the starting values is not essential. In this way, we scan a range of *r*_0_ and *δ_ρ_* and use iBME to fit *I*(0), *cst*, and the conformational ensemble. To simplify interpretations and analysis, we kept the parameter *θ* constant for each protein (values specified in Table 1). This was done to keep the balance between the prior and experiment constant, so as to focus on changes that arise due to differences in the hydration and displaced solvent parameters. The resulting reweighted ensembles (at different values of *r*_0_ and *δ_ρ_*) are analyzed further below.

**Table 1:** Best fitting SAXS parameters, input and results of the iBME optimization

|  | Hst5 | Sic1 | Tau | TIA-1 |
| --- | --- | --- | --- | --- |
| $r_0$ [Å] | 1.722 | 1.558 | 1.640 | 1.722 |
| $\delta\rho$ [$e/nm^3$] | 10.02 | 10.02 | 0.00 | -3.34 |
| $\theta$ | 80 | 80 | 50 | 100 |
| $\chi^2_{red}$ (before iBME) | 3.52 | 1.39 | 1.64 | 0.919 |
| $\chi^2_{red}$ (after iBME) | 1.04 | 1.02 | 1.14 | 0.540 |
| $\phi_{eff}$ | 0.911 | 0.941 | 0.707 | 0.884 |

We also validated the grid scanning approach using synthetic SAXS data generated using a specific choice of *r*_0_ and *δ_ρ_* to generate the data (see SI text). The results show that both with a correct prior (Fig. S6) and a prior that is different from that used to generate the synthetic data (Fig. S7) the method is able to recover the values of *r*_0_ and *δ_ρ_* that were used to generate the data.

### A scoring function for the ensembles on the *r*_0_ × *δ_ρ_* grid

Once we calculated SAXS profiles and refined (i.e. reweighted) the ensembles for each pair of parameters on the *r*_0_ × *δ_ρ_* grid, we needed a scoring function to quantify the agreement with the experimental data. We already note here the complication arising from the fact that the ensembles have been refined against the experiments.

We first calculated the 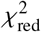 after the iBME optimization to indicate the quality of the ensembles. For Hst5, Sic1, Tau and TIA-1 this lead to a large region with low-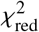 (Fig. 1a-d-g-j), suggesting that most of the combinations of *r*_0_ and *δ_ρ_* tested with *r*_0_ ≤ 1.722 Å can be fitted to the experimental data. We note also that although the 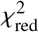 is widely used for the purpose of comparing SAXS profiles, it has been noticed that it can be prone to overfitting if the noise is not estimated correctly (74, 75). For this reason, previous studies have focused e.g. on identifying the amount of information in a SAXS profile or in correcting the experimental noise (75–77). As we are here comparing different fits to the same data and with the same number of degrees of freedom, we did not use such corrections here. In turn this means that the calculated values of 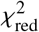 cannot easily be compared across the four systems that we analyzed.

**Figure 1:**
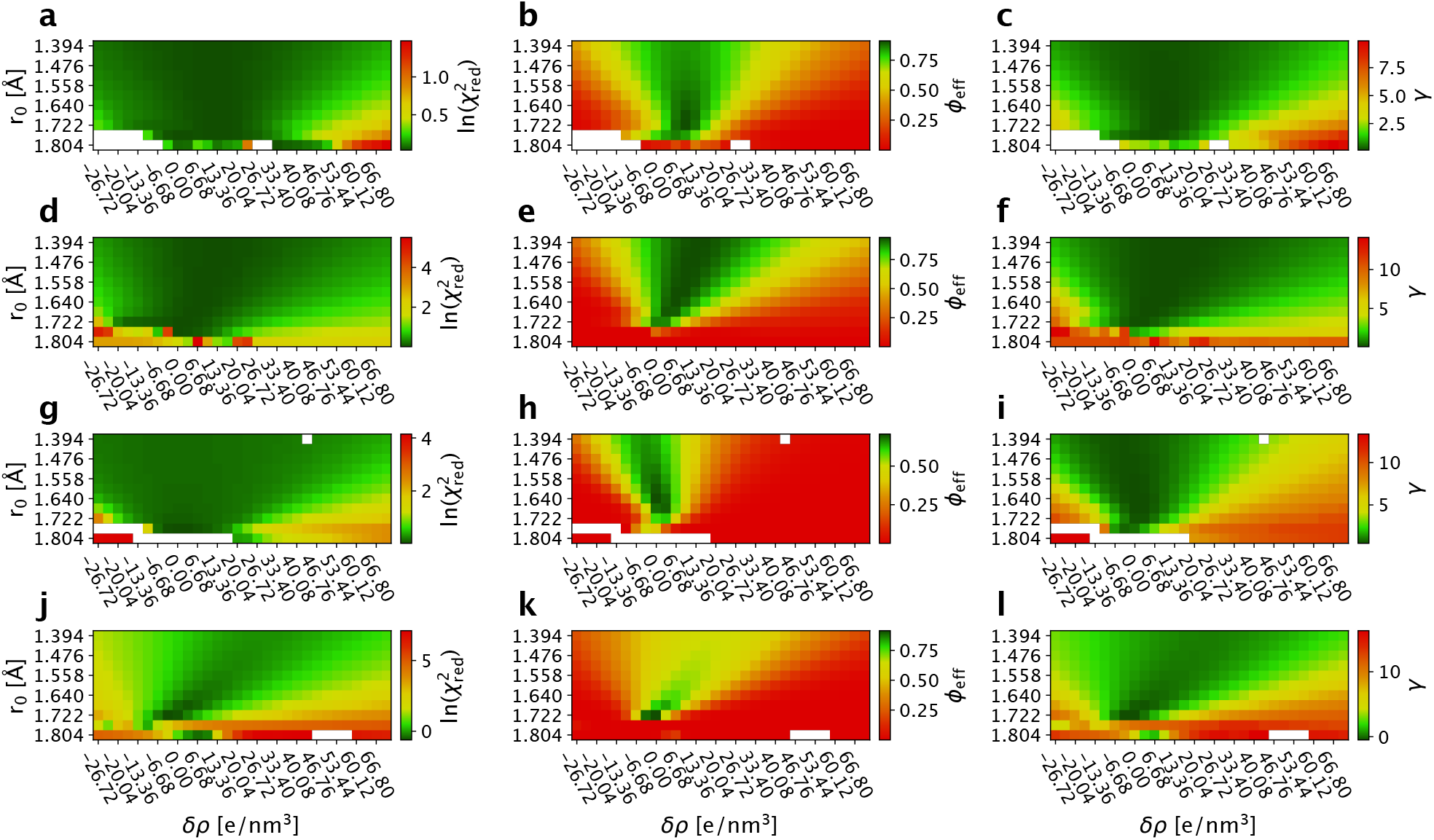
Reweighting ensembles using SAXS data calculated using different values for the parameters that effect the contribution from for the hydration layer and displaced solvent. The grids show the results from the iBME ensemble optimization with different combinations of *δ_ρ_* and *r*_0_. The top row (a–c) shows Hst5, the second row (d–f) shows Sic1, the third row (g–i) shows Tau, and the last row (j–l) shows results for TIA1. For each protein we show in the first column (a, d, g, j) ln 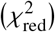, the second column (b, e, h, k) *ϕ*_eff_, and third column (c, f, i, l) 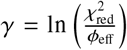. White spots correspond to ensembles where the iBME reweighting failed.

When reweighting an ensemble against experiments, it is important to monitor the effective fraction of frames (*ϕ*_eff_) that, as explained above, quantifies how much the posterior distribution deviates from the prior. When *ϕ*_eff_ is low this indicates that the ensemble has to be modified substantially to achieve the desired agreement with experiments. In order to ease comparison across the different ensembles in the grid, we have chosen to use the same value of *θ* for all combinations of *r*_0_ and *δ_ρ_*, where *θ* sets the balance between not deviating too much from the prior ensemble (maximizing *ϕ*_eff_) and fitting the experimental data (minimizing 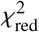). Thus at fixed values of *θ* the resulting value of *ϕ*_eff_ is another indicator of the quality of the ensembles (78), and we find a relative narrow region of the grids with high values of *ϕ*_eff_ (Fig. 1b,e,h,k). Thus comparing the maps of 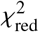 and *ϕ*_eff_ we find that while it is possible to achieve a relative good fit at a wider range of values of *δ_ρ_* and *r*_0_, in many cases this comes at the cost of a substantial deviation from the prior (low *ϕ*_eff_). To combine the balance of achieving a low 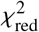 and a high *ϕ*_eff_ we thus introduce a variable, 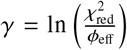, that combines these two effects in a single number (Fig. 1c,f,i,l). The results show that it is not possible to obtain a good fit (defined here as giving rise to a low *γ*) at all values of *δ_ρ_* and *r*_0_, but that there are certain regions that appear to give rise to comparable fits. The parameter sets that gives the lowest values of *γ* for Hst5, Sic1, Tau and TIA-1 are reported in Table 1 together with the 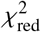 before and after reweighting and the *ϕ*_eff_. We note that the final ensemble may not be optimal, and that further refinement could be obtained by scanning *θ* (24, 36, 37).

We observe that while there are some differences between the four proteins analyzed above, it also appears that there is a region that gives relative good fits for all proteins (Fig. 1). Because it may be computationally expensive to scan many sets of parameters, we also aimed to find a set of parameters that provides good scores for these four proteins. We therefore normalised and averaged the *γ* scores, and found the minimum to be at *δ_ρ_* = 3.34 *e/nm*^3^ and *r*_0_ = 1.681 Å.

We note that in the literature higher values are generally reported as default for *δ_ρ_* (generally 10% (7) or 6% (17, 72) of the bulk density). In the context of SAXS calculations, however, we also note that the main quantity that determines the contribution of the hydration layer is the product between *δ_ρ_* and its width Δ (13, 14). Whereas Δ is 3 Å in Crysol, it is chosen in a slightly different fashion in Pepsi-SAXS and it is 5 Å in most of the cases that we examined. To demonstrate that different values for the contrast of the hydration layer alone can lead to the same result when the width is treated in different ways, we also repeated the grid scans for Hst5, Sic1, Tau and TIA-1 employing FoXS. Although the minima for *γ* are different from those obtained using Pepsi-SAXS (Fig. S8), the reweighted distributions of *R_g_* from these minima are essentially identical (Fig. S9), reinforcing the observation that *δ_ρ_* alone is meaningful only in the context of a specific SAXS calculator. In addition, by again normalizing and averaging the *γ* score, we obtain a global minimum for the parameters in FoXS at *r*_0_ = 1.68 Å (as in Pepsi-SAXS) and *δ_ρ_* = −7.07 *e/nm*^3^. We note that this value for *δ_ρ_* appears substantially different from those used for folded proteins, and suggest that further studies are needed to examine better the physical origins of these effects.

Conversely, the value *r*_0_ = 1.681 Å is slightly higher than the average values used by Crysol and FoXS (1.62 Å) and Pepsi-SAXS (1.64 Å). While the origin of this observation is unclear, we note that there are differences in protein volume depending on whether a protein is folded or unfolded (79, 80). Thus, the excluded volume—as described by the Fraser model (59)—might need different parameters for compact and expanded proteins, and we suggest that this could be studied further using molecular simulations (9).

### Effect of hydration and atomic radius parameters on the conformational ensemble

The idea of the grid search is to find a combination of *r*_0_ and *δ_ρ_* that gives rise to the best agreement with experimental data, taking also into account that we need to determine the parameters and ensemble weights at the same time. Here we explore the effect of choosing specific sets of these parameters on the conformational ensembles.

We first examined how much the individual ensembles differed from that determined using the *r*_0_ and *δ_ρ_* parameters that give rise to the lowest value of *γ* (Table 1). We therefore calculated, as a measure of the difference between ensembles, *ϕ*_eff_ between the weights optimized using the different combinations of *r*_0_ and *δ_ρ_* relative to the weights obtained using the ‘optimal’ values of *r*_0_ and *δ_ρ_* (Fig. 2). As expected, values around the optimum give rise to comparable weights (*ϕ*_eff_ close to 1). For Sic1 and TIA-1 we also note a correlation between *r*_0_ and *δ_ρ_*, so that increasing the excess density (*δ_ρ_*) and decreasing the radius (*r*_0_) appears to give rise to more comparable ensembles. Nevertheless, the results also show that while several different combinations of *r*_0_ and *δ_ρ_* can give rise to a good fit (Fig. 1), the resulting ensembles differ depending on the choice of parameters used to calculate the scattering data. In particular, we find that the ensembles are rather sensitive to the choice of *δ_ρ_*, in particular for the three IDPs analyzed above.

**Figure 2:**
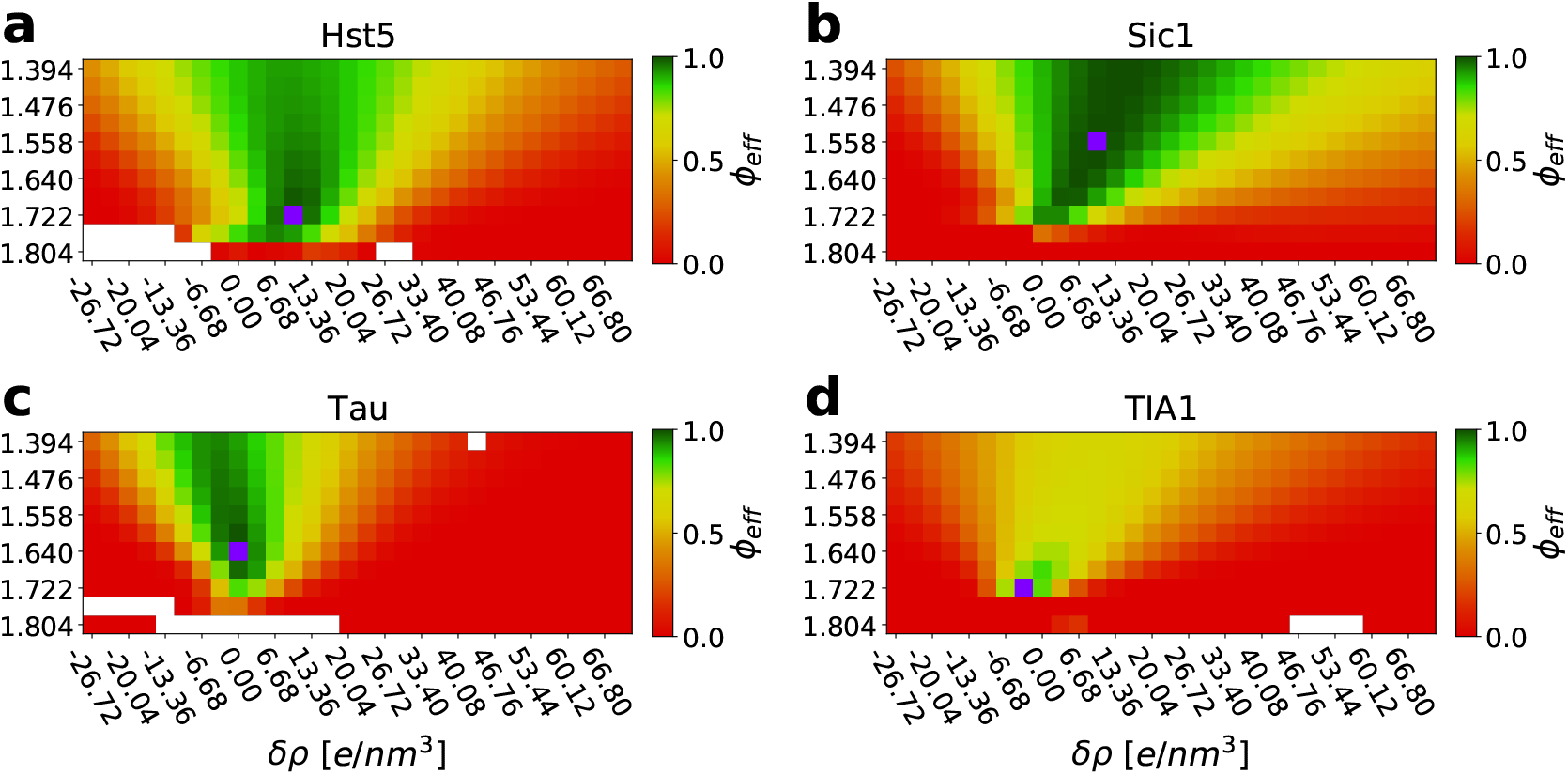
Comparing ensembles relative to the optimum. For each protein we calculated the effective fraction of frames (shown here as *ϕ*_eff_) between the weights obtained using the parameters in Table 1 and the weights obtained at all other combinations of *r*_0_ and *δ_ρ_*. White spots correspond to ensembles where the BME reweighting failed. Purple spots correspond to the minima for *γ*.

SAXS data are often used to estimate the radius of gyration (*R_g_*), and so we demonstrate how the different ensembles have different distributions of *R_g_*. Using Sic1 as an example we chose the optimal parameters as well as three other combinations of *r*_0_ and *δ_ρ_* and calculated *p*(*R_g_*) after reweighting (Fig. 3). The results show that as long as *r*_0_ and *δ_ρ_* are chosen within the range that gives a low value of *γ* the resulting distribution is relatively similar. On the other hand, if more extreme values for the *r*_0_ and *δ_ρ_* parameters are chosen, the average *R_g_* may differ substantially in the reweighted ensembles (Fig. S10).

**Figure 3:**
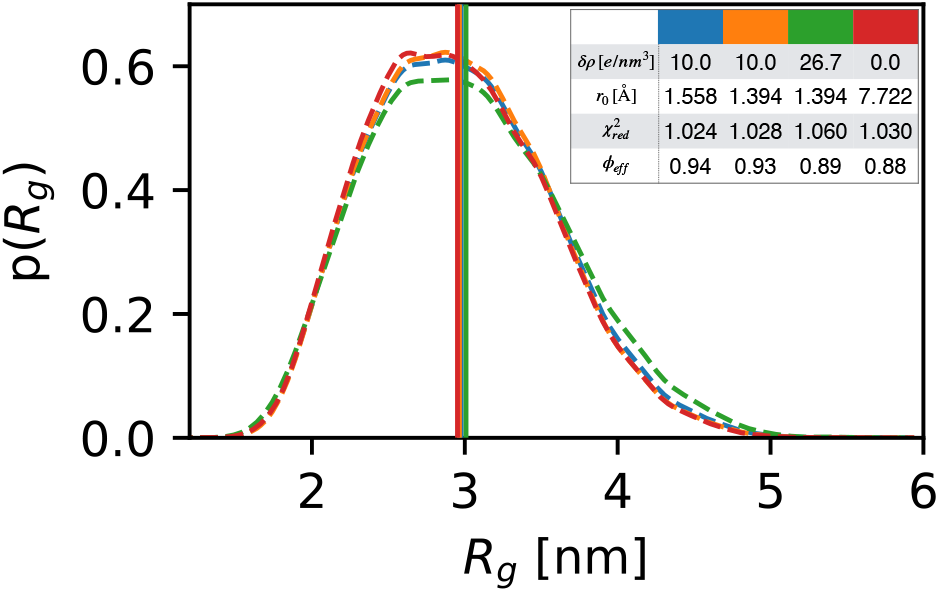
Effect of the *δ_ρ_* and *r*_0_ parameters on reweighted probability distributions of *R_g_*. We use Sic1 as an example and show *p* (*R_g_*) from both the optimal (lowest *γ*) parameters (blue) as well as three other choices of *r*_0_ and *δ_ρ_* in the low-*γ* region (orange, green and red). The insert shows the parameters used in each case and the results of the reweighting on the *R_g_* distribution.

### Assessing the influence of the prior on the parameters search

As our strategy to determine self-consistent values for *δ_ρ_* and *r*_0_ is based on the Bayesian/Maximum Entropy refinement of probability distributions, it is reasonable to ask how the results depend on the statistical prior used in the approach. In this context, there are two related questions that we here address. First, as we have also examined previously (66, 68), is the question of how much the distribution of conformations after reweighting depends on the prior that is used. Second, is the question of how much the *δ_ρ_* and *r*_0_ parameters depend on the prior. The latter is important because the optimal parameters may in part compensate for imperfections in the prior.

To examine these questions we applied our protocol to three different ensembles of *α*-Synuclein. The first ensemble was generated using Flexible-meccano, while the other two were previously generated by molecular dynamics simulations (67) using either the Amber a99SB-*disp* (disp) or the Amber ff03ws (a03ws) force field. The distributions of the *R_g_* for the three priors are relatively different (Fig. 4a) and, consequently, the minima of the *γ* parameter indicate small differences in the best values of *δ_ρ_* and *r*_0_ (Fig. 5). The reweighted distributions of *R_g_*, however, appear very similar (Fig. 4b). Notably, for each prior we obtain essentially indistinguishable distributions of *R_g_* whether we use the optimal parameters (for each prior) from the grid search or the set of parameters that we proposed as default values (Fig. 4b).

**Figure 4:**
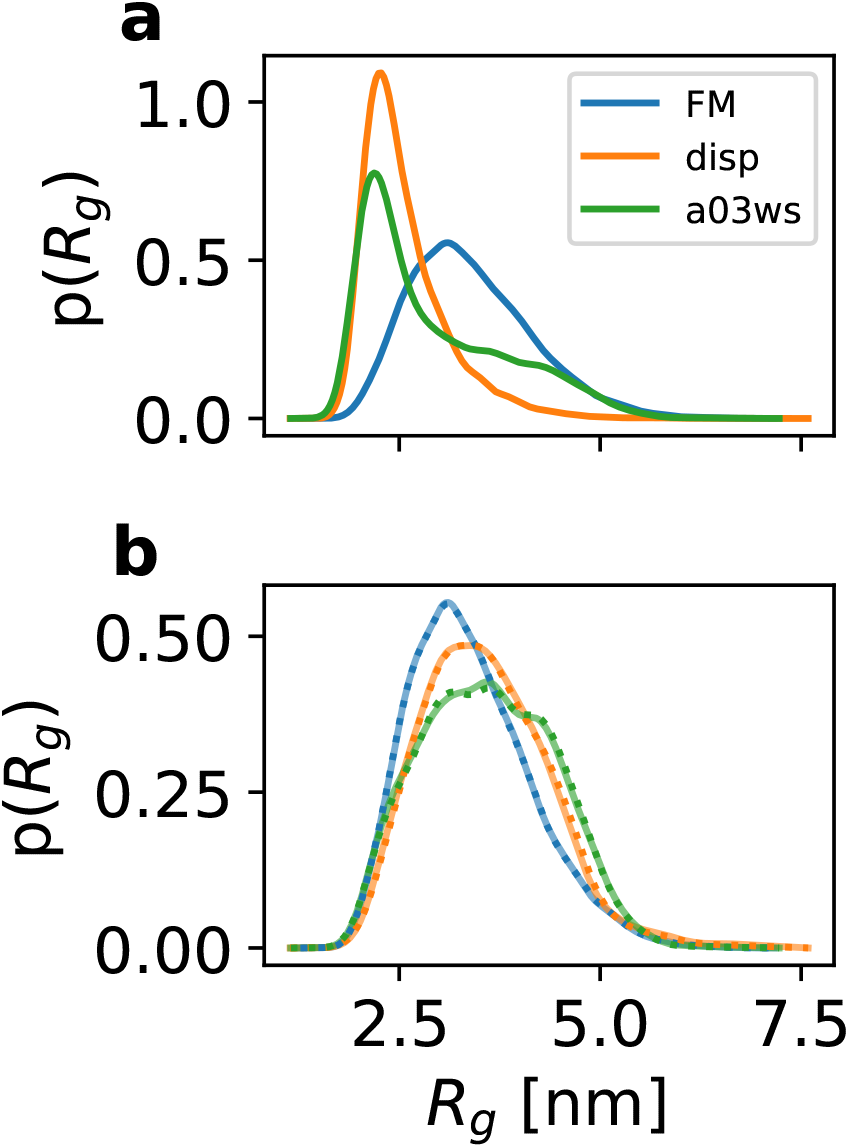
Effect of the prior distribution. (a) Distributions of *R_g_* of *α*-Synuclein sampled with Flexible-meccano (FM), a99SB-*disp* (disp) and a03ws. (b) Reweighted *R_g_* distributions either from the optimal (lowest *γ*) *δ_ρ_* and *r*_0_ parameters for each ensemble (solid lines) or using the default values we propose (*δ_ρ_* = 3.34 *e/nm*^2^ and *r*_0_ = 1.681 Å; dotted lines).

**Figure 5:**
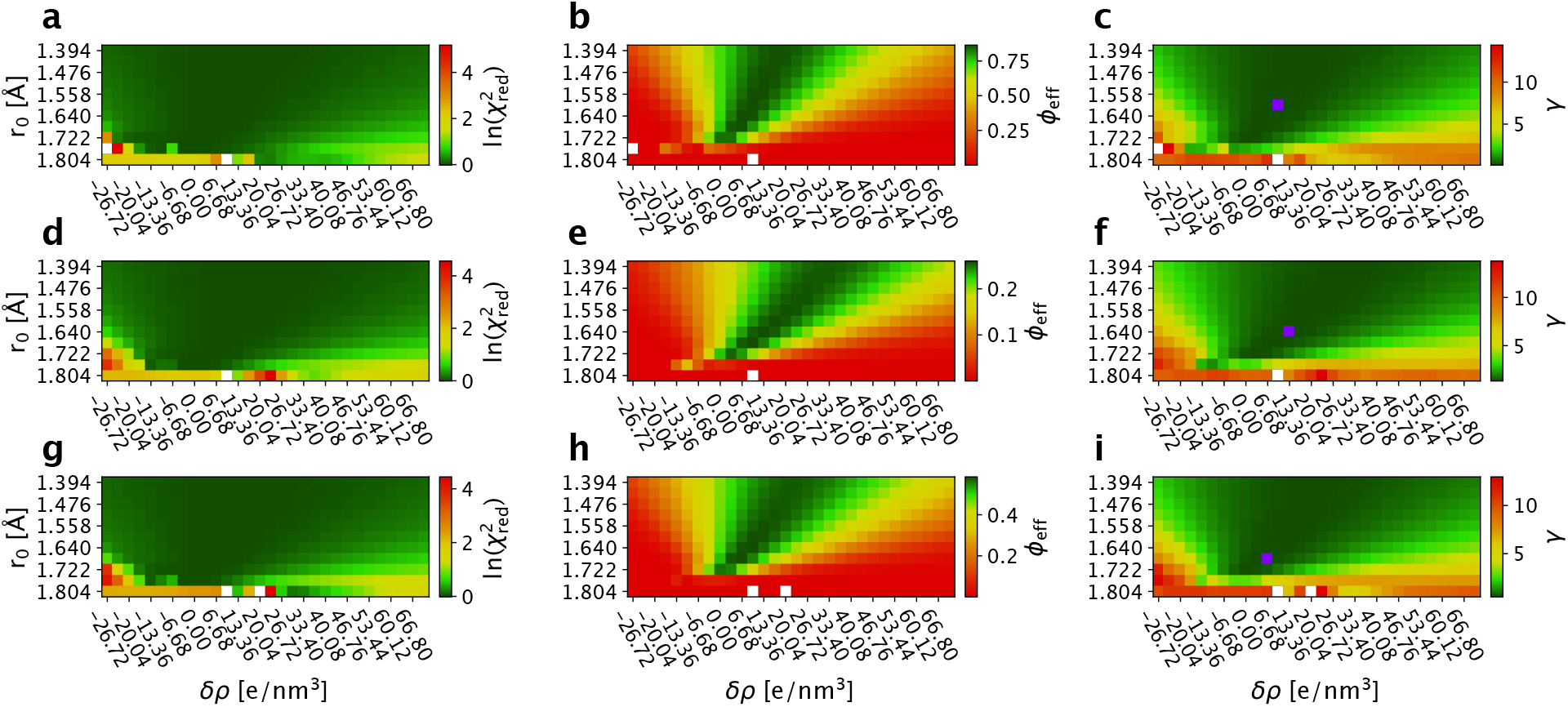
Reweighting *α*-Synuclein ensembles using SAXS data calculated using different values for the parameters that effect the contribution from for the hydration layer and displaced solvent. The grids show the results from the iBME ensemble optimization with different combinations of *δ_ρ_* and *r_0_*. The top row (a–c) shows the results from the Flexible-meccano ensemble, the second row (d–f) shows the results using a99SB-disp as the prior, and the third row (g–i) shows the results from a03ws as the prior. For each ensemble we show in the first column (a, d, g) ln 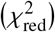, the second column (b, e, h) *ϕ*_eff_, and third column (c, f, i) 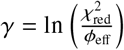. White spots correspond to ensembles where the iBME reweighting failed. Purple spots in the third column correspond to the minima for *γ*.

Returning to the original two questions, these results show that the prior may influence the optimal parameters resulting from the grid search, similar to our observations using synthetic SAXS data (Figs. S6 and S7). They also show, in line with previous observations (66, 68), that even when starting from somewhat different priors, the posterior distributions tend to be substantially similar. Noteworthy, the results are robust to the choice of *δ_ρ_* and *r*_0_ so that very similar results are obtained even when using the global minimum from our analyses of Hst5, Sic1, Tau and TIA-1.

### Comparing ensembles to experimental estimates of *R_g_*

In the analyses of the *R_g_* described above, we implicitly referred to the values calculated from the protein coordinates as the mass-weighted root-mean-square distance from the centre of mass of the protein. This is a geometric quantity that is often used to study protein behaviour and biophysics. Because the ensembles were constructed by fitting to the experimental SAXS data, the resulting averages and distributions of *R_g_* represent the experimental system, but exactly because the hydration effects were included in the SAXS calculations, this means that these *R_g_* values only represent the protein.

Another approach to estimate the average *R_g_* (〈*R_g_*〉) from experiment is to fit the SAXS data directly without resorting to a conformational ensemble. The most common approach is to use the Guinier approximation (60), though other approaches exist (81–83). Because the SAXS data potentially contain a contribution from the hydration layer, the 〈*R_g_*〉 estimated by a Guinier analysis (or similar methods) may in principle contain contributions from this (17). One complication of a Guinier analysis is to identify the linear part of the curve (the Guinier region), in particular because the first few low-*q* points of the scattering curve may often be noisy. As rule of thumb, the maximum scattering angle that can be used for the Guinier approximation satisfies the condition *q*_max_〈*R_g_*〉 < 1.3 (1), but a threshold value of 0.9 has also been proposed for disordered systems (84).

Because both approaches to estimate 〈*R_g_*〉 are commonly used, we here compare the two results. In addition to shedding light on differences, this analysis is also relevant because it is relatively common to compare 〈*R_g_*〉-values calculated from simulations with values estimated from experiments, even though the two might differ due to effects of the hydration layer. We thus performed a Guinier analysis of the SAXS data for the four proteins, progressively extending the upper limit of the *q*-range from 0.9 to 1.3 and plotting *R_g_* vs *q*_max_ *R_g_* (84). We find that the Guinier fits can show substantial differences in the estimated 〈*R_g_*〉 values depending on the range used. Returning to the question of how the *R_g_* values estimated from the Guinier fit compares to the average *R_g_* from the conformational ensembles with the lowest *γ* scores (horizontal black line in Fig. 6), we find that these are in a reasonable agreement (within 0.2 nm) with the values calculated from Guinier fits using *q*_max_ *R_g_* = 1.3. Looking across the four proteins we do not find a unique *q*_max_ *R_g_* value where the Guinier fit gives rise to an average *R_g_* that is similar to that obtained from the conformational ensembles.

**Figure 6:**
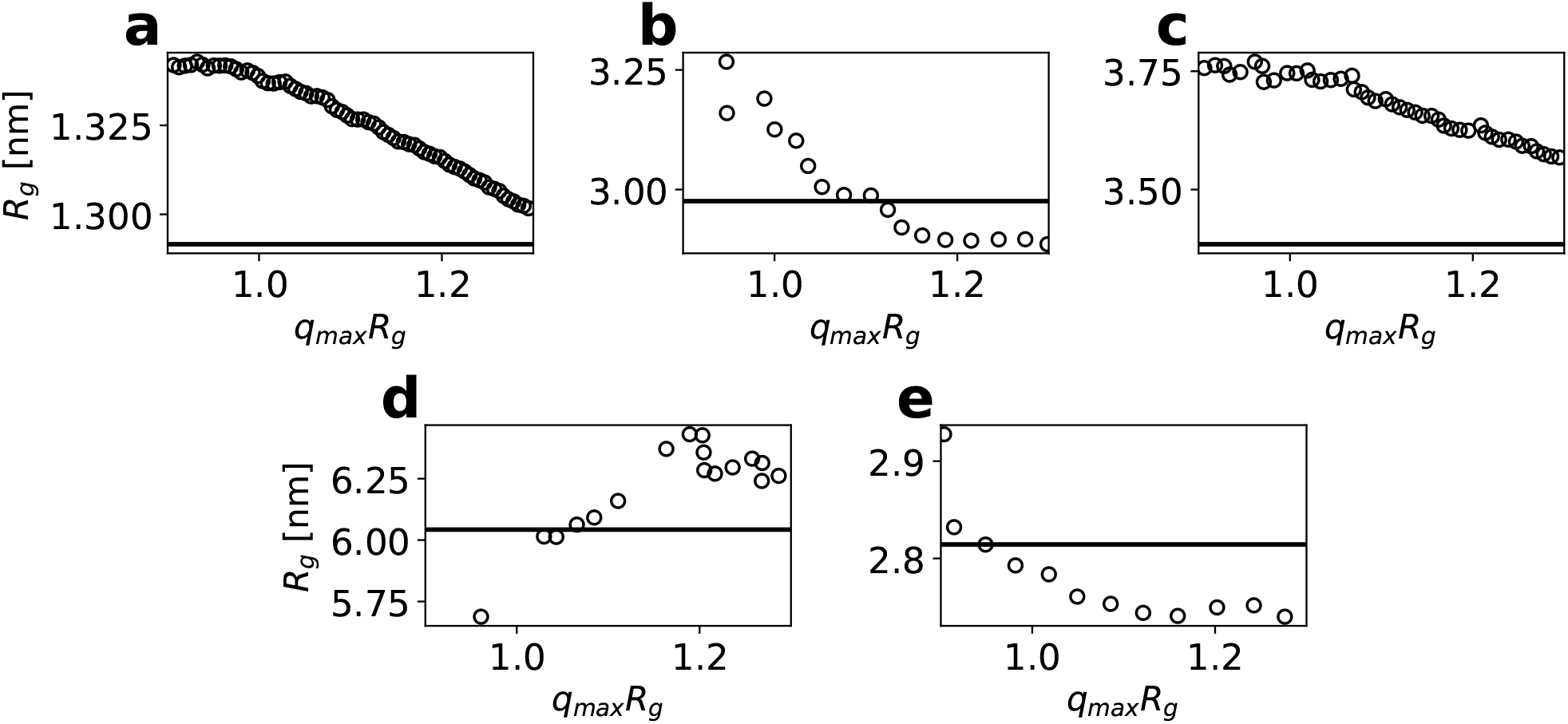
Estimating 〈*R_g_*〉 from experimental SAXS profiles of (a) Hst5, (b) Sic1, (c) Tau and (d) TIA-1 using Guinier fitting and ensemble refinement. We used the Guinier approximation to estimate *R_g_* by fitting from the lowest measured value of *q* (in the case of Hst5 we ignored the first 10 points due to noise) to different values of *q*_max_, reporting the results as *R_g_* vs. *q*_max_*R_g_* (black circles). The horizontal black lines are the ensemble averaged *R_g_* calculated from the conformational ensembles with the chosen optimal *r*_0_ and *δ_ρ_* parameters (Table 1).

## CONCLUSIONS

SAXS experiments are widely used as source of structural information and are often integrated with computational methods to determine conformational ensembles. Generally, such approaches rely on a forward model such as Pepsi-SAXS (14) to calculate SAXS data from one or more conformations and optimizes the structures or weights to improve agreement with experiments. While these approaches are very powerful, they are subject to uncertainty due to the choice of unknown parameters in the forward model. In principle these parameters can be ‘integrated out’ using Bayesian approaches (31, 85), though this can become computationally prohibitive for SAXS calculations. The aim of our work is thus to provide a robust protocol that estimates values for the relevant free parameters. In the context of SAXS these include the two parameters that determine the effects of the hydration layer and displaced volume (*δ_ρ_* and *r*_0_), as well as a scale factor and constant background (*I*(0) and *cst*) that are often necessary to estimate.

We have developed and tested iBME as an extension to the weight estimation in BME to include a scale factor and constant background between experimental and calculated values. Importantly, the values are estimated as the globally best fitting parameters and are determined self-consistently with the weights of the ensembles. While we have here presented iBME in the context of SAXS data, other types of data, such as NMR residual dipolar couplings, solvent paramagnetic relaxation enhancement effects or circular dichroism spectra, may also involve estimating an overall scale. For small-angle neutron scattering data, the ability to include (fit) a constant background can be important because of contributions from incoherent scattering.

We also present the results from an extensive analysis of the effect of the *r*_0_ and *δ_ρ_* parameters on calculated SAXS data and the resulting ensembles. We have determined self-consistent ensembles where the ensembles are reweighted using SAXS data calculated using different values for these parameters. Such an analysis is in particular important for large ensembles of flexible molecules, since fitting these parameters to each conformation could lead to substantial overfitting. We also note that the calculations of SAXS intensities could potentially be improved further by being able to predict the features and contribution from the hydration layer for different sequences and conformations, rather than relying on fitting parameters.

Combining these two aspects, the approach that we have described can be summarized as follows: (i) Sampling a conformational ensemble. (ii) Calculate SAXS profiles from the conformers of the ensemble, keeping scale and background parameters fixed (*I*(0) = 1, *cst* = 0), and perform a grid scan for *δ_ρ_* and *r*_0_. (iii) For each value of *δ_ρ_* and *r*_0_, optimize the weights, *I*(0) and *cst* using iBME. (iv) Examine the results by calculating 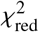, *ϕ*_eff_ and *γ*, and select the ensemble with the lowest value of *γ*.

One complication of the algorithm is that it requires a large number of calculations of SAXS intensities. In cases where the prior already exists or is fast to generate, the SAXS calculations can quickly become limiting in terms of computational efficiency. For these reasons we also propose default values for *δ_ρ_* and *r*_0_ that we find to provide relatively accurate results for the four proteins that we examined. To test this further, we also used these default parameters to calculate SAXS intensities from different conformational ensembles of *α*-Synuclein and find that the resulting distributions of *R_g_* are almost the same as if the parameters are optimized. We also note that the computational overhead of the grid scans could be drastically reduced by precomputing partial SAXS intensities once per grid, and then adding the contributions from *δ_ρ_* and *r*_0_. Although this procedure is already internally used by several methods to calculate SAXS data, options to output and process partial intensities for specific scattering angles are, at the moment, not easily accessible.

Finally, we also discuss considerations on the common practice of comparing the experimentally determined *R_g_* (calculated with the Guinier approximation) with *R_g_* calculated from the structural ensemble. While the results show good agreement, they also suggest that caution should be exerted when comparing average *R_g_* values from simulations and experiments. In particular, we find that the both changing the *r*_0_ and *δ_ρ_* parameters (Fig. S10) or the region used for Guinier fitting (Fig. 6), can change the *R_g_* substantially, and so generally we recommend that it is better to compare the experimental data (in this case SAXS intensities) with values calculated from simulations, rather than comparing parameters estimated from experiments. Nevertheless, even such comparisons contain ambiguities as one needs to chose parameters in the SAXS calculations. We thus suggest that our work will be useful when benchmarking molecular simulations against SAXS data by providing additional insight into the effect of the hydration layer (17) and suggest default values that can be used as a starting point.

## ACKNOWLEDGMENTS

We acknowledge Sandro Bottaro for fruitful discussions and for implementing the iBME algorithm in BME, Simone Orioli for useful discussions and input about SAXS and other aspects of this manuscript. We thank Andreas Haahr Larsen and Ramon Crehuet for discussions and comments on the manuscript. We are grateful to Dimitri Svergun and Tanja Mittag for sharing SAXS data for, respectively, Tau and Sic1. This research was funded by the Lundbeck Foundation BRAINSTRUC initiative in structural biology (to KLL; R155-2015-2666).

## Notes

https://github.com/KULL-Centre/papers/tree/main/2021/SAXS-pesce-et-al

https://github.com/KULL-Centre/BME

### Competing Interest Statement

The authors have declared no competing interest.

### Summary of Updates

New analyses and updated text

