## Supporting Text, Figures and Table for "Refining conformational ensembles of flexible proteins against small-angle X-ray scattering data"

#### 1 Validation of iBME with synthetic SAXS data

We used the Hst5 ensembles generated by Flexible-Meccano to create synthetic data to test and validate iBME. We calculated SAXS profiles for each structure in the ensemble using Pepsi-SAXS, using $\delta\rho = 13.36\ e/nm^3$ and $r_0 = 1.722$ Å as parameters to describe the hydration layer and displaced solvent volumes. These SAXS profiles were linearly averaged to give rise to the target SAXS data, assigning the error associated with the $j$'th data point as $\sigma_j = \frac{0.5 I_j}{100} \exp(q_j)$

We then generated five sets of non-uniform weights for the ensemble, to define five different prior distributions. Specifically, we generated weights for each conformer, $i$, according to:

$$w'_i = \exp\{1 - (20 + 10a) * \exp[(0.4b + 1.2) - R_g(i))^4]\} \tag{1}$$

with $a$ and $b$ being random numbers between 0 and 1, and with the final weights ($w_i$) obtained by normalizing $w'_i$. These weights lead to the SAXS data in Fig. S1) and $R_g$ distributions shown in Fig. S2.

First we use the standard BME approach with $\theta = 100$ to optimize each of the five priors against the (synthetic) experimental data by minimizing the functional

$$\mathcal{L}(\omega_1 \cdots \omega_n) = \frac{m}{2}\chi^2_{\text{red}}(\omega_1 \cdots \omega_n) - \theta S_{\text{rel}}(\omega_1 \cdots \omega_n) \tag{2}$$

as described in the main text and with $\chi^2_{\text{red}}$ defined as

$$\chi^2_{\text{red}}(\omega_1 \cdots \omega_n) = \frac{1}{m}\sum_i^m \frac{(\sum_j^n \omega_j F(x_j) - F_i^{EXP})^2}{\sigma_i^2} \tag{3}$$

After doing so we keep the resulting weights ($\omega_j$) as reference.

Standard BME assumes that the experimental observable ($F^{EXP}$) are on the same scale as those obtained by applying the forward model to the computational ensemble ($F(x_j)$). iBME is an approach that deals with cases where the two are on a different scale, such as for example SAXS data where $I^{EXP}$ and the calculated $I(x_j)$ may differ by a linear transformation $I'(x_j) = \text{scale} \cdot I(x_j) + \text{offset}$.

To generate synthetic data representing this situation, we thus changed each of the input SAXS curves (for each of the structures) by multiplying by a random number between 0 and 5 to change the scale, and subsequently adding a random number between 0 and 1 to change the offset (the same scale and offset was used for each structure in the ensemble). The average curves are shown in Fig. S3.

We then applied iBME (as described in the main text) to these priors, targeting the unmodified synthetic data (blue line in Figs. S1 and S3). The same $\theta$ as with standard BME (100) was used. The successful outcome of iBME is demonstrated by comparing the $\chi^2$ both before and after reweighting, $\phi_{\text{eff}}$ and and the weights obtained using BME on the unmodified SAXS data (Fig. S4).

### 2 Validation of the grid scan with synthetic SAXS data

We use the same synthetic experimental SAXS data used to test iBME above (i.e. to fit the scale and offset) to assess the ability of the grid scan procedure to recover the $\delta\rho$ and $r_0$ used to generate the synthetic experimental SAXS profile.

We first used uniform weights (same as used to generate the synthetic experimental SAXS profile) as the prior for the iBME optimization. Even when adding noise to the (synthetic) data, the grid search recovers the correct values used to generate the synthetic data ($\delta\rho = 13.36 e/nm^3$ and $r_0 = 1.722$Å) at the minimum of $\gamma$ (determined by a $\chi^2_{red} \approx 0$ and $\phi_{eff} \approx 1$) (Fig. S6).

We also repeated the grid scan using 'Prior 1' (Figs. S2–S3) discussed above as the prior. In this way, we represent the case where the experimental data are generated by a different distribution than the prior. In this case we find the minimum of $\gamma$ in a grid point adjacent to that used to generate the synthetic experimental SAXS profile ($\delta\rho = 10.02\ e/nm^3$ and $r_0 = 1.763$Å) (Fig. S7). Thus, while we do not recover exactly the same values, they are very close to those used to generate the data.

### 3 Additional figures and tables

Table S1: Search ranges for fitting parameters

| | CRYSOL | FoXS | Pepsi-SAXS |
|---|---|---|---|
| $r_0$ [Å] | 1.55 - 1.68 | 1.40 - 1.80 | 1.56 - 1.72 |
| $\delta\rho$ [$e\ nm^{-3}$] | 0 - 70.0 | -27.0 - 54.0 | 0 - 33.4 |

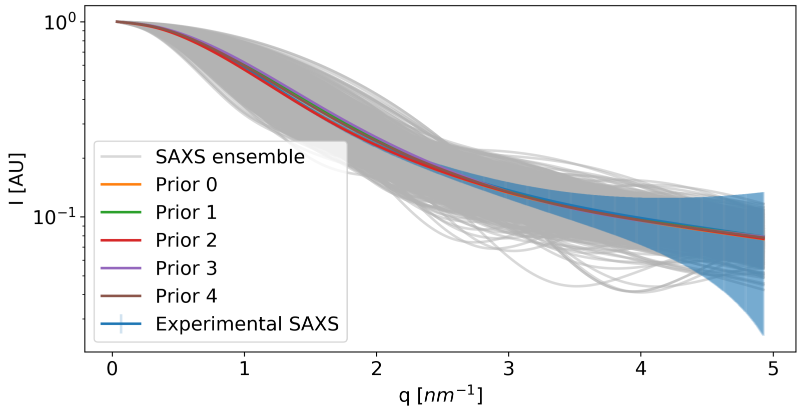


Figure S1: Synthetic SAXS data used to validate the iBME protocol. Thin grey lines show SAXS profiles for each of the structures in the Hst5 ensemble. The uniform average of these curves gives rise to the blue line, which we here term the (synthetic) 'experimental' data. This is the target for the optimization. The non-uniform weights give rise to five other average SAXS curves, that are the starting point for optimization.

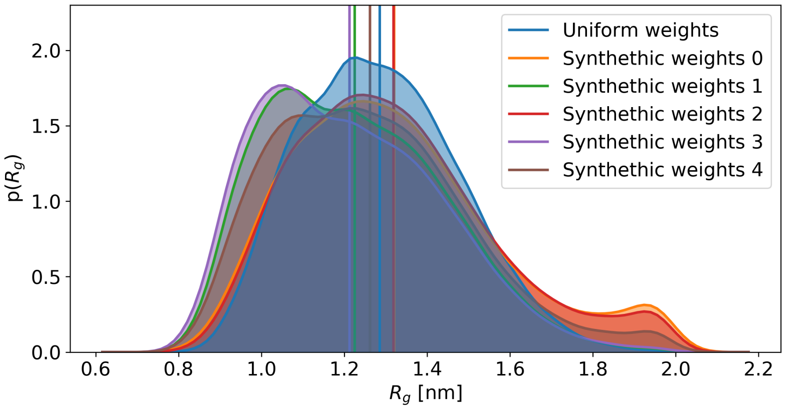


Figure S2: $R_g$ distribution from the Flexible-meccano ensemble of Hst5 (with uniform weights), as well as five ensembles with different sets of non-uniform weights.

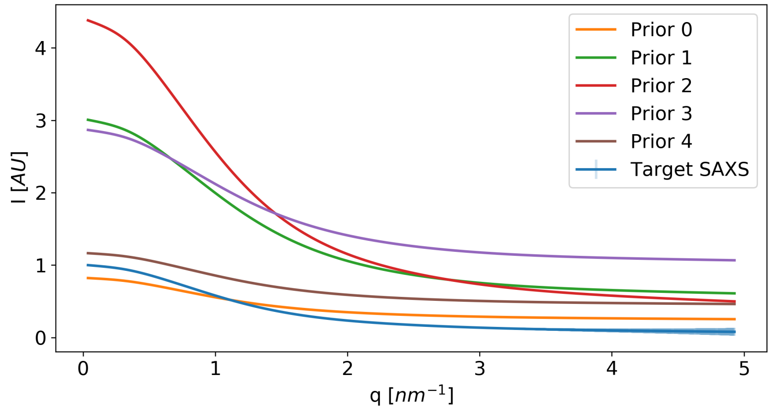


Figure S3: Same SAXS profiles as in Fig. S1, but after perturbations with a scale and offset.

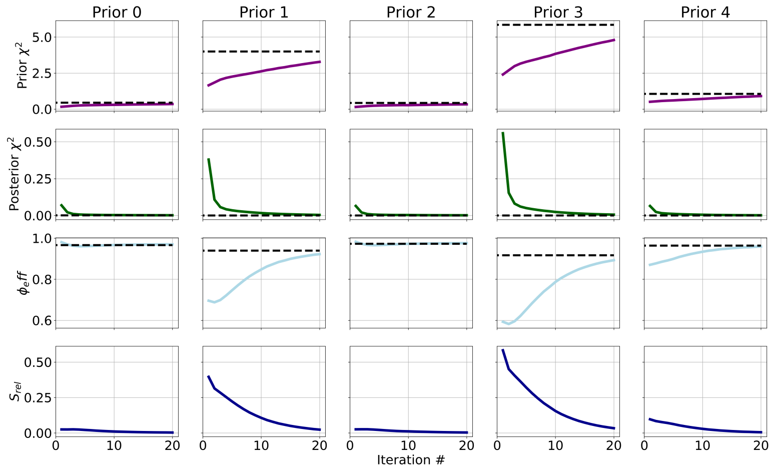


Figure S4: Evolution of observables along the iterations of the iBME. Dotted black lines represent the target values obtained from the standard BME using the un-scaled and shifted data. The relative entropy $S_{rel}$ is computed between the weights at each iteration of iBME and those obtained from standard BME.

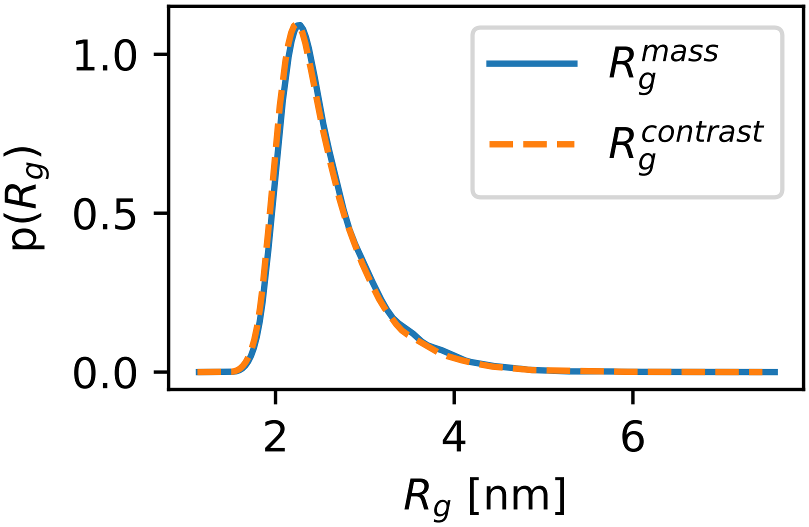


Figure S5: Distribution of the mass-weighted $R_g$ and contrast-weighted $R_g$ calculated as described in the Methods of the main text for the a99SB-disp ensemble of $\alpha$-Synuclein.

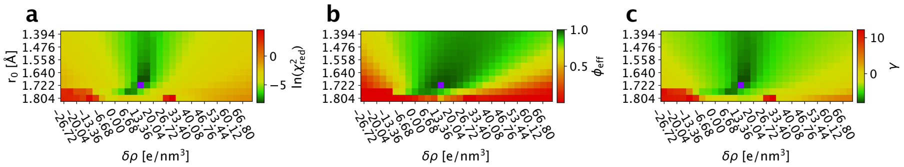


Figure S6: Grid scan optimizing a synthetic experimental SAXS profile with iBME. In this case we used as prior the same distribution as that used to generate the synthetic data. The minima in $\chi^2_{red}$ (a), $\phi_{eff}$ (b) and $\gamma$ are shown in purple.

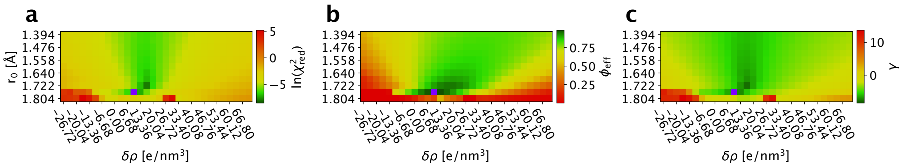


Figure S7: Grid scan optimizing a synthetic experimental SAXS profile with iBME. As prior for iBME we use 'Prior 1' (Figs. S2 and S3). Minima in $\chi^2_{red}$ (a), $\phi_{eff}$ (b) and $\gamma$ are shown in purple.

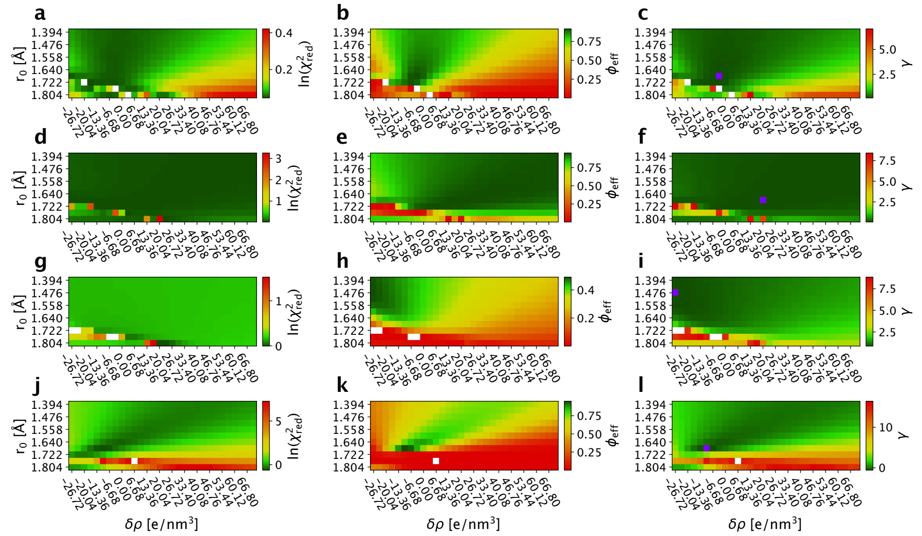


Figure S8: Reweighting ensembles using SAXS data calculated with FoXS using different values for the parameters that effect the contribution from for the hydration layer and displaced solvent. The grids show the results from the iBME ensemble optimization with different combinations of $\delta\rho$ and $r_0$. The top row (a–c) shows Hst5, the second row (d–f) shows Sic1, the third row (g–i) shows Tau, and the last row (j–l) shows results for TIA1. For each protein we show in the first column (a, d, g, j) $\ln\left(\chi^2_{\mathrm{red}}\right)$, the second column (b, e, h, k) $\phi_{\mathrm{eff}}$, and third column (c, f, i, l) $\gamma = \ln\left(\frac{\chi^2_{\mathrm{red}}}{\phi_{\mathrm{eff}}}\right)$. White spots correspond to ensembles where the iBME reweighting failed. The purple spots in the third column correspond to the minima for $\gamma$.

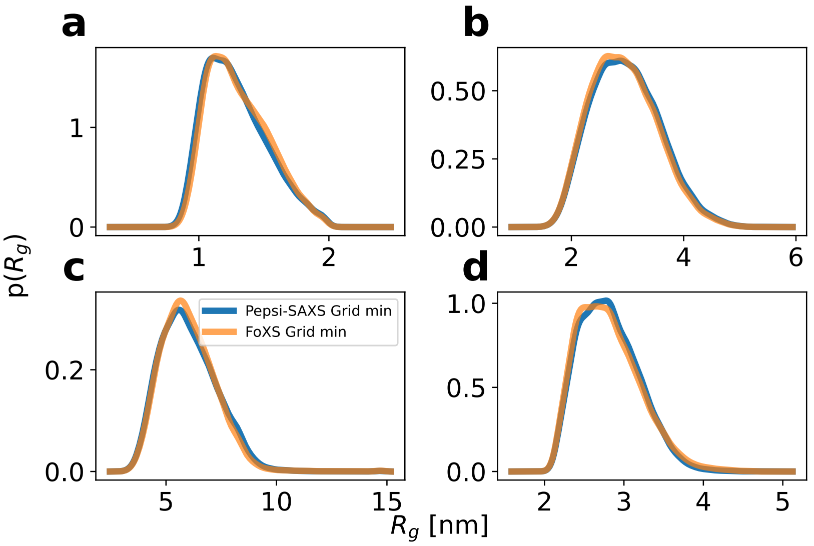


Figure S9: Reweighted $R_g$ distributions for (a) Hst5, (b) Sic1, (c) Tau and (d) TIA-1 from the $\gamma$ minima obtained with either Pepsi-SAXS or FoXS-based grid scans.

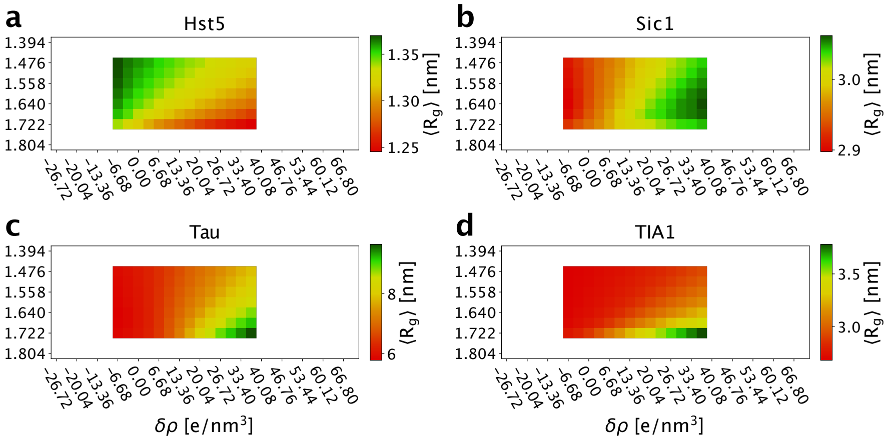


Figure S10: Reweighted average values of $R_g$ on the part of the grids that gave reasonable fits for (a) Hst5, (b) Sic1, (c) Tau and (d) TIA-1.
